# *Staphylococcus aureus nt*5 gene contributes to bacterial infection ability to form kidney abscess

**DOI:** 10.1101/2022.02.09.479838

**Authors:** Yang Ye, Xinpeng Liu, Zijing Xia, Min Tang, Dan Zhang, Fuqiang He, Peng Zhang, Shufang Liang

**Affiliations:** State Key Laboratory of Biotherapy and Cancer Center, West China Hospital, Sichuan University, and Collaborative Innovation Center for Biotherapy, No.17, 3rd Section of People’s South Road, Chengdu, 610041, P. R. China; Department of Rheumatology, West China Hospital, Sichuan University, Chengdu, 610041, Sichuan, P. R. China; Department of Urinary Surgery, West China Hospital, Sichuan University, Chengdu, 610041, Sichuan, P. R. China

**Keywords:** *Staphylococcus aureus*, *nt*5 gene, 5’-nucleotidase, gene silence, abscess, phagocytosis, daptomycin

## Abstract

*Staphylococcus aureus* (*S. aureus*) is a commonly conditional infection pathogen, in which several key virulence genes are responsible for bacterial infection ability. The *S. aureus nt*5 gene, encoding 5’-nucleotidase, mediates bacterial nucleic acid pathway, yet it is nearly unknown of *nt*5 function for staphylococcal infection ability. Herein we have constructed *S. aureus* mutant with the gene *nt*5^C166T^ silence (*S. aureus* Δ*nt*5) by a CRISPR RNA-guided base editing system to investigate bacterial infection ability *in vitro* and *in vivo*. As expected, several *nt*5-related genes are disturbed in *S. aureus* Δ*nt*5, in which gene transcription level of *py* is decreased compared with the wild-type *S. aureus*. Bacterial gene *nt*5 is downregulated and *py/adk* are upregulated when *S. aureus* is exposed to antibiotics daptomycin, which indicates *nt*5-mediated nucleic acid pathway is interfered upon with daptomycin treatment. Furthermore, the mutant Δ*nt*5 displays about a 40-fold reduction of bacterial loading in mouse kidney on a mouse sepsis model, and the infection ability of Δ*nt*5 is reduced than the wild-type bacteria. The gene *nt5* contributes to *S. aureus*-infected abscess formation in mouse kidney, and the silence of *nt5* gene promotes phagocytosis of *S. aureus* by mouse and human immunocytes. In general, our findings reveal *nt5* silence impedes bacterial loading in kidney to form abscess but enhances *S. aureus* to be phagocytosed by host cell immune system *in vitro* and *in vivo*, which indicates that *nt5* gene plays an important role in bacterial infection and immune evasion.

## Introduction

*Staphylococcus aureus* (*S. aureus*) is a commonly conditional pathogen causing severe infections in almost every organ of human body, including sepsis, endocarditis, skin and wound infections, or osteomyelitis (1). Treatment of *S. aureus*-induced infections has become increasingly challenging due to emergence of drug-resistant bacteria (2). Drug resistance to many clinically used antibiotics becomes more and more serious, even several powerful antibacterial antibiotics, such as (DAP), vancomycin and linezolid, have suffered from antibiotics resistance (3,4). Emergence of drug resistance always overspeeds the development of new antibiotics. Hence, it is necessary to explore key pathogenicity-driven genes and their infection mechanism of *S. aureus*, which contributes to develop novel therapeutic means against *S. aureus* infections.

DAP is a novel cyclic lipopeptide antibiotic, which interacts with bacterial cell membrane to efficiently kill most clinically relevant Gram-positive bacteria (5). Our previous research has identified several key DAP-targeting bacterial proteins through quantitative drug proteomic analysis on the conditional pathogenic bacteria *S. aureus* (6). The downregulation of *nt*5 (gi 446956624) upon DAP exposure leads to decrease of bacterial membrane potential and disruption of cell membrane, which contributes to bacterial DNA rapid release and finally causes bacteria death (6). Both genes *py* (gi 487015361) and *adk* (gi 488431505) are essential in *nt*5-mediated adenine metabolism of *S. aureus* (7). DAP disrupts kinase activity of *py-*encoding pyruvate kinase (PY, protein ID: ODV52833.1) and ADK (protein ID: WP_002500890.1) to mediate adenine metabolism pathway (6). These evidences indicate bacterial *nt*5 gene and bacterial adenine synthesis are tightly related with DAP antibacterial effect.

The *nt*5-encoding 5’-nucleotidase (NT5, protein ID: ABD29455.1) protein is ubiquitous among bacteria species, including *E coli, S. aureus* and *Streptococcus suis* (8), which hydrolyzes the phosphate group of 5′-nucleotide. Most of the soluble intracellular NT5 from bacteria belongs to the vast haloacid dehalogenase superfamily, such as *E. coli* NagD (9) and *S. aureus* NT5. In *Bacillus subtilis*, YcsE and YktC as major NT5s both exhibit 5′-nucleotide phosphatase activity (10). *Streptococcus suis* NT5 is an important virulence factor enabling to inhibit neutrophil functions *in vitro* by converting adenosine monophosphate to adenosine (Ado) (8). More noticeably, bacterial NT5 activity is associated with immune evasion properties (11). But it is nearly unclear how does *S. aureus* NT5 function in *S. aureus*-induced infection before.

During infection, Staphylococcal nuclease and adenosine synthase A (adsA, protein ID: ALY19673.1) converts neutrophil extracellular traps to deoxyadenosine (dAdo), which is toxic to human immune cells and kills phagocytes and macrophages (12). The Ado and dAdo suppress immune responses and have immunoregulatory attributions (13,14). In *S. aureus*, adsA encoded by gene *ads*A (gi 59698588) is a cell wall-anchored enzyme to convert AMP/dAMP to Ado/dAdo, contributing to *S. aureus* escaping from phagocytic clearance and formation of organ abscesses *in vivo* (15). The adsA harbors NT5 signature sequence which is critical for catalytic activity of mammalian enzymes (16). NT5 activity of *S. aureus* is a mediator of virulence attributes of adsA (15). Although NT5s are widespread among all domains of life, it is unclear whether *S. aureu* NT5 with a 33.3kD-molecular weight has similar properties as adsA.

To assess the role of NT5 activity toward DAP treatment and Staphylococcal infection ability, herein we focus on gene function of *S. aureus nt*5 and its effects on bacterial infection ability. Using lose-of-functional mutant strain with *nt*5^C166T^ gene silence performed by a CRISPR RNA-guided base editing system, we have revealed *S. aureus nt*5 gene is required for *S. aureus*-induced abscess formation. Moreover, we have confirmed the antibiotics DAP exposure to *S. aureus* leads to downregulation of genes *nt*5 and *adsA*, and upregulation of genes *py* and *adk*, therefore *nt*5-mediated bacterial nucleic acid metabolic pathway is disturbed by DAP. We also have revealed the silence of *nt*5 gene promotes the killing of *S. aureus* by immune cells. Our findings indicate disruption of the *nt*5-invovling bacterial activities provides a potentially novel therapeutic means against *S. aureus* even the emerging antibiotic-resistant pathogens.

## Materials and methods

### Reagents and chemicals

DAP with 99% purity was bought from Dalian Meilun Biotechnology Company, China. Kanamycin and chloramphenicol were ordered from Sigma-Aldrich for plasmid screening in *E. coli* and plasmid curing in *S. aureus* RN4220 and *S. aureus* ATCC 25923. *Bsa*I-HF and T4 DNA ligase were purchased from New England Biolabs for Golden gate assembly. The iScript cDNA Synthesis Kit and SsoAdvanced SYBR GreenSupermix were bought from Bio-Rad for qRT-PCR.

Human lymphocyte separation solution (DKW-LSH-0100) was ordered from Beijing Dakewei Company. MojoSort human CD14^+^ monocytes isolation kit was purchased from Biolegend. Cell culture media DMEM and RPMI-1640 were purchased from Hyclone.

### Strains and plasmids

The bacterial strains including *S. aureus* ATCC 25923 and *S. aureus* RN4220 were stored with 20% glycerol (7). *S. aureus* was streaked from glycerol stocks to culture on tryptic soy broth (TSB), then was inoculated into 5 mL TSB to grow overnight at 37 °C with constant shaking for usage. *E. coli* was grown in Luria–Bertani broth.

The base editing plasmid pnCasSA–BEC was endowed from Professor Ji of ShanghaiTech University. The pnCasSA–BEC system mainly includes the APOBEC1-nCas9 that is a fusion protein composed of a deaminase APOBEC1 and a Cas9 nickase Cas9D10A, cap1A and rpsL promoters and a replication origin ColE1 (17). In addition, the plasmid contains an antibiotic kanamycin selection marker for plasmid amplification in *E. coli*, and another chloramphenicol-resistance marker for plasmid selection in *S. aureus*. The repF is a temperature-sensitive origin for plasmid curing in *S. aureus*.

### Plasmid construction of pnCasSA-BEC-NT5sp

We designed a 20 bp-spacer oligo to insert in *Bsa*I site of the pnCasSA-BEC plasmid by Golden Gate assembly. The plasmid from the Golden Gate assembly reaction was transformed into 100 μL competent *E. coli* DH5α cells to incubate at 37°C overnight, and the recombinant plasmid pnCasSA-BEC-NT5sp was sequenced for verifying correctness of plasmid construction.

### *S. aureus* competent cells

The electrocompetent cells of *S. aureus* strains were prepared as following procedures. A single colony of each *S. aureus* strain was inoculated into 2 mL TSB to incubate at 30°C for 12 h. Cells were diluted with a ratio of 1:100 into 100 mL fresh TSB medium, then were cultured up to OD_600_ value with 0.3–0.4 and chilled on ice for 20 min. Cells were harvested by centrifugation with 5000 rpm at 4°C for 5 min, and washed twice with 0.5M sterile ice-cold sucrose. Cells were suspended into 1 mL 0.5M sucrose, from which 100 uL aliquots were frozen in liquid nitrogen and stored at -80°C [18].

### Modified plasmid transformation by electroporation

The pnCasSA-BEC-NT5sp plasmid was transformed into the competent cells of laboratory strain *S. aureus* RN4220 by electroporation to get the modified plasmid. Before cell electroporation, 100uL competent cells and the plasmid pnCasSA-BEC-NT5sp were respectively thawed on ice for 5 min. The cells were mixed with 1 ug of the recombinant plasmid, and transferred into an electroporation cuvette with 1 mm-thickness at room temperature. After being pulsed at 2.1kV mm, 100U and 25mF, cells were immediately supplied with 900μL TSB and recovered at 30°C for 1.5h. Finally, 500μL cells were plated onto a TSB plate containing 5 mg/L chloramphenicol for incubation at 30°C for 24 h (18). The genomic DNAs of the target colonies were extracted to confirm base mutation in the target site by sequencing.

### Base editing in *S. aureus* 25923

To acquire *nt*5^C166T^ gene silence (Δ*nt*5) in the conditionally infectious pathogen *S. aureus* ATCC 25923 (*S. aureus* Δ*nt*5), the assembled plasmid isolated from *S. aureus* RN4220 was transformed into *S. aureus* ATCC 25923 by electroporation performed as above. 1 mL cells were plated onto a TSB plate containing 5 mg/L chloramphenicol to incubate at 30 °C for 24 h. The target colonies were screened to extract genomic DNAs for sequencing to confirm *nt*5^C166T^ in *S. aureus* ATCC 25923. The method of plasmid curing was referenced Ji’s article (18).

### Primer design for gene mutation and qRT-PCR

We selected a 20 bp-spacer sequence before a PAM site (NGG) in the target gene locus for designing sgRNAs. Two oligos were designed as qRT-PCR primers (Table 1). The total RNA was isolated using bacterial RNA extraction kit following the manufacturer’s protocol (RE-03111, FOREGENE Company). The expression levels of genes *nt*5 and *py, adk* were normalized to that of the reference *Tpi* gene. Each gene expression was quantified by qPCR for three times (n=3).

### Minimal inhibitory concentration (MIC) of DAP

The MIC of DAP on *S. aureus* 25923/Δ*nt*5 was determined by broth microdilution following the Clinical Laboratory Standards Institute guidelines. DAP was serially diluted to final concentrations ranged from 0.03125 to 16 µg /mL. The MIC was measured by three experiment repeats.

### Mouse sepsis model

The following experiments on BALB/c mice, including a mouse sepsis model, immune evasion and phagocytosis, were approved by the Animal Care Committee of Sichuan University (Chengdu, China). The infection ability of *S. aureus* ATCC 25923 (WT) and Δ*nt*5 was respectively assayed on a mouse sepsis model. Overnight cultures of each *S. aureus*, including the WT and Δ*nt*5, were diluted 1:100 into fresh TSB and grown for 3 h at 37°C. Then each *staphylococci* culture was centrifuged, washed twice, and diluted in PBS to yield an OD_600_ up to 0.5 with 10^8^ colony-forming units (CFU)/mL. 10^6^ CFU bacterial suspension was administered via tail intravenous into a 6-week-old male BALB/c mouse. Totally ten mice were inoculated with each strain culture, however, two mice died of *S. aureus* WT infection during experiment progression.

On the 5^th^ day of bacteria inoculation, mice were killed to observe the abscess formation in kidneys and staphylococcal survival in kidneys and livers. Mouse’s left kidney and left lobe liver were respectively collected to homogenize with PBS containing 1% Triton X-100. Aliquots of homogenates were diluted and plated on TSB agar plates for triplicate measurements of CFUs. Histopathology analysis of mouse right kidney tissue was performed as following. Tissues were incubated at room temperature in 10% formalin for 24 h, and embedded in paraffin to be sectioned, stained with hematoxylin-eosin to examine by microscopy. Data were obtained from each representative group samples with cohorts of six mice.

### Cell culture

The murine macrophage RAW 264.7 cells and human myeloid leukemia mononuclear THP-1 cells were respectively cultured in Dulbecco’s Modified EagleMedium (DMEM, Hyclone) and Roswell Park Memorial Institute (RPMI 1640, Hyclone) media supplemented with 10% fetal bovine serum (FBS) at 37 °C and 5% CO_2_. Cells were cultured for 48h, and collected to adjust cell density to 10^7^ cells/mL for assay.

### Phagocytosis to *S. aureus* 25923/Δ*nt*5 *in vitro*

*S. aureus* 25923 WT and mutant Δ*nt*5 were cultured overnight, from which 1 mL bacteria was cultured with 100 mL TSB medium for additional 3 h at 37°C to detect their phagocytosis by monocytes. As mentioned above, each staphylococci culture was centrifuged, washed twice, and diluted in PBS to yield an OD600 up to 0.5 with 10^8^ colony-forming units (CFU) /mL. 10^6^ CFU bacterial suspension was mixed with 5×10^5^ RAW264.7 or THP-1, and cultured at 37°C from 15 to 90 min respectively to plate on TSA and count on the next day.

The following experiments were approved by the West China Hospital Medical Center Institutional Review Board of Sichuan University, following the guidelines of the West China Hospital Medical Center Institutional Review Board of Sichuan University. The enrolled volunteers provided written consent to participate in this study. To further measure *S. aureus* 25923/Δ*nt*5 survival ability under the existence of *ex vivo* neutrophils, 1mL peripheral blood was respectively collected from 8 BALB/c mice and 5mL peripheral blood was collected from 3 human volunteers. 2.5×10^4^ CFU bacteria was mixed with 225 µL of mouse blood, and 10^6^ CFU bacteria was mixed with 900 µL of human blood to incubate at 37°C with slow rotation [15]. The mixture of bacteria with mouse or human blood were sampled after incubation for 30 and 90 minutes, from which aliquots were incubated with 0.5% saponin/PBS on ice for 30 min. Dilutions of staphylococci were plated on TSA for enumeration of surviving CFUs.

### Isolation of human serum CD14^+^ monocyte to test phagocytosis

The remaining 4ml peripheral blood samples were collected from each donor and diluted with isotonic PBS to isolate lymphocyte using separation solution (DKW-LSH-0100, Beijing Dakewei Company). After centrifugation with 250g for 10 min, cells in the plasma layer were collected to isolate CD14^+^ monocytes using a commercial kit (MojoSort human CD14^+^ monocytes isolation kit, 480047, Biolegend) according to the manufacturer’s protocol. Generally, the cells were resuspended with MojoSort™ buffer to adjust to 10^8^ cells, from which 100 uL suspension was taken into a new centrifuge tube to mix with 10 uL Biotin-antibody cocktail and 5 uL Human TruStain FcX™ to incubate at room temperature for 10 min. Then 10 uL Streptavidin nanobeads were mixed to incubate on ice for 15 min, and cells were cleaned and collected by centrifugation at 300g for 5 min, then 2.5 mL of Mojosort™ was added to collect the decanted liquid on a centrifuge tube via a magnet device. Finally, 5×10^5^ cells were mixed with 10^6^ CFU bacteria, cultured at 37°C, sampled at 30, 60 and 90 min respectively, plated and counted the survival rate of bacteria.

### Immune evasion ability of *S. aureus* 25923 /Δ*nt*5 detected on mouse

In order to detect the immune evasion ability of *S. aureus* 25923/Δ*nt*5 *in vivo*, we injected 10^6^ CFU bacteria of *S. aureus* 25923/Δ*nt*5 from the tail vein for 5 BALB/c mice, and collected 100 µL blood from mouse orbit after infection for 30, 90 min. Aliquots were incubated with 0.5% saponin/PBS on ice for 30 min to lyse host eukaryotic cells. Dilutions were plated on TSA plates for enumeration of surviving CFUs.

### Hemolysis assay *in vitro*

*S. aureus* WT and Δ*nt5* were grown in TSB at 37 °C overnight and diluted in TSB to yield an OD600 up to 0.5 with 10^8^. Then 1 μL of culture of each strain including *S. aureus* 25923 and Δ*nt*5 was loaded onto a sheep blood plate [17] to incubate at 37 °C for 24 h for observing their hemolytic circle size.

## Results

### Construction of *nt*5^C166T^ mutant by a CRISPR RNA-guided editing system

To investigate the loss-of-function of *nt*5 gene, we constructed gene silence mutant strain Δ*nt*5 by a base editing CRISPR RNA-guided cytidine deaminase system pnCasSA–BEC which includes APOBEC1-nCas9, cap1A and rpsL promoters [17]. In pnCasSA-BEC system, the deaminase is directed to any genomic locus by the Cas9/sgRNA complex and directly catalyzes the conversion of cytidine (C) to uridine (U) through a deamination reaction. This base editor has an editable window from positions 4 to 8 in the spacer sequence and it can achieve C/T conversion without homologous recombination (Fig. 1A), which is efficient for gene inactivation in *S. aureus*.

**Fig. 1.**
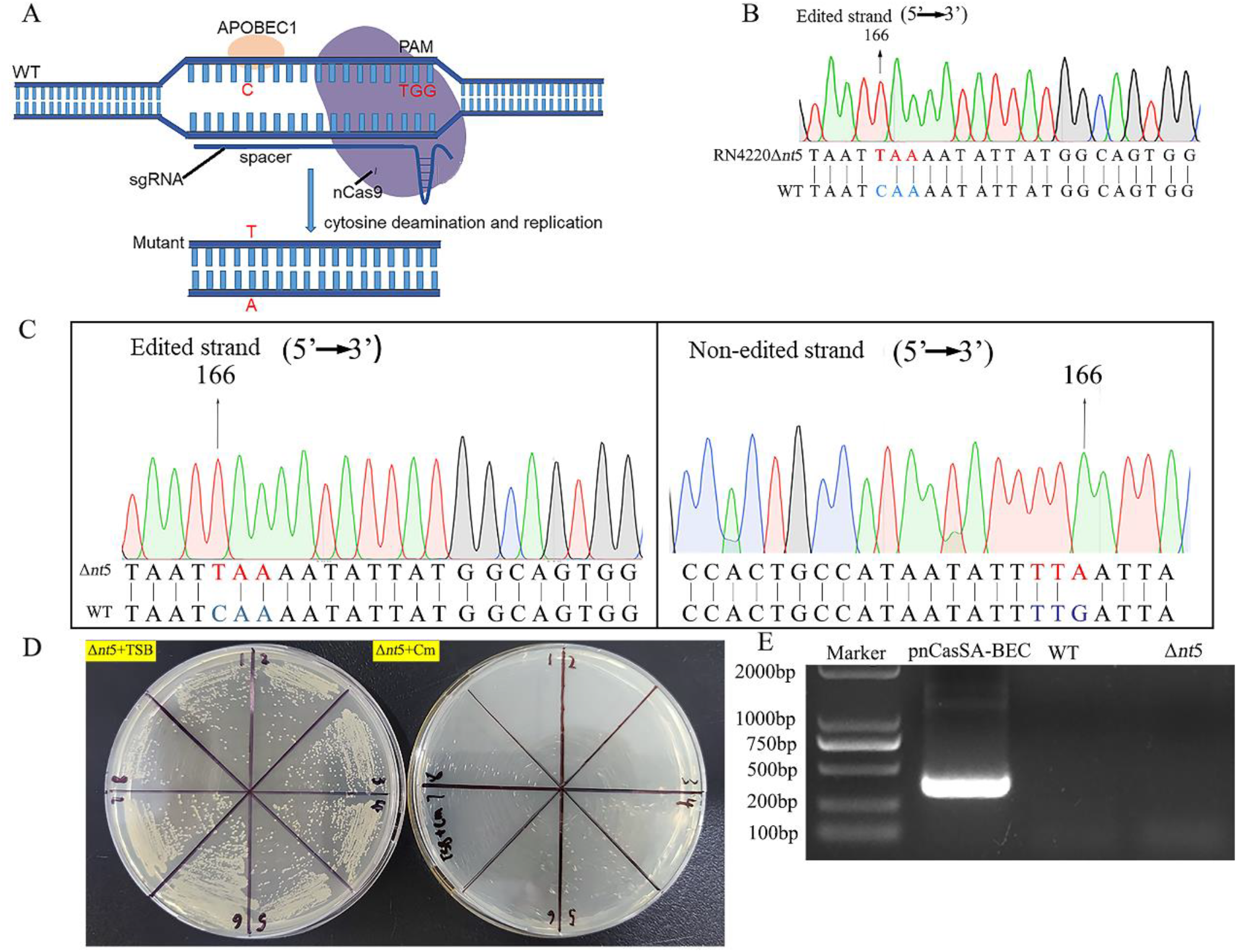
Construction of *nt5*^C166T^ gene silence *S. aureus* Δ*nt5*. (A) Composition of the base editor pnCasSA-BEC system. It mainly includes APOBEC1-nCas9, cap1A and rpsL promoters, and it can be directed to any genomic locus by the Cas9/sgRNA complex to achieve C/T conversion in positions 4 to 8 of the spacer sequence without homologous recombination. PAM: protospacer adjacent motif. (B-C) The C166 of *nt5* gene was mutated to T in the edited nucleotide strand in the *S. aureus* RN4220 (B) and *S. aureus* 25923 (C) respectively. (D-E) The pnCasSA-BEC-NT5sp plasmid was easily cured from *S. aureus* Δ*nt5* after base editing. The individual colonies obtained after the curing steps were streaked onto the TSB agar plates in the presence (D, right) or absence (D, left) of Cm to test the effectiveness of curing (D), and (E) the PCR result showed that the assembled plasmids disappeared in Δ*nt5*. Cm: chloramphenicol, WT: wild type.

We constructed the pnCasSA-BEC-NT5sp plasmid by Golden Gate assembly, then each of 200 ng/μL plasmids was transformed into the laboratory strain *S. aureus* RN4220 with 100% efficiency by electroporation. The base C166 of *nt*5 gene in *S. aureus* RN4220 was mutated to T (*nt*5^C166T^), which led to produce a premature stop codon with TAA (Fig. 1B). Correspondingly, the mutant *nt*5-encoding protein 5’-nucleotidase (mNT5) was a truncated mutant in which the natural amino acid Q56 of the wild-type NT5 was mutated and terminated due to emergence of the premature stop codon TAA.

*S. aureus* RN4220 retains the methylation function of the *Sau* I system, which thus appropriately methylates DNA for subsequent uptake by other target *S. aureus* strains containing restriction–modification systems. Firstly, we transferred the recombinant plasmids pnCasSA-BEC-NT5sp modified by *S. aureus* RN4220 to our target object *S. aureus* ATCC 25923, in which the pnCasSA-BEC-NT5sp system enabled highly efficient C/T conversion for target base editing. The C at position 7 of spacer in *nt*5 gene was mutated to T with 100% efficiency (Fig. 1C), which indicated that C166 of *nt*5 gene of the *S. aureus* ATCC 25923 strain was mutated to T so that Q56 amino acid of the wild-type NT5 protein was interrupted and terminated at translational level due to emergence of the mutant premature stop codon TAA.

Although the gene *nt*5 was not deleted, the endogenous encoding wild-type protein NT5 was interrupted to lose their normal functions. After plasmid curing, four randomly picked colonies grew normally in the absence of chloramphenicol whereas none of them could grow in the presence of the antibiotic after two passages (Fig. 1D). Meanwhile, no assembled plasmid was detectable in *S. aureus* Δ*nt*5 (Fig. 1E), which confirmed a stable removal of the pnCasSA-BEC-NT5sp plasmid from the host strain *S. aureus* ATCC 25923. All base mutations were verified by sequencing.

### The *nt*5 mutation leads to expression decrease of gene *py* and *adk*

The gene *adk*, and *py* are essential in *nt*5-mediated nucleic acid metabolism of *S. aureus* based on the reported literature and bioinformatics analysis by STRING software (http://string-db.org/cgi/input.pl) (19). We further verified if *nt*5 gene mutation had influences on the two gene transcription levels. Therefore, the gene transcription levels of *nt5, adk and py* were detected on mutant Δ*nt5* to learn about their gene level changes along with the single base mutation of gene *nt5*.

As results, compared with WT, the RNA levels of *nt*5 *and py* in *S. aureus* Δ*nt*5 were significantly decreased by 2.82 and 1.90 times respectively (p<0.05, p<0.01) (Fig. 2A and 2B). Gene transcriptional level of *adk* was 1.55-times of downregulation, which indicated the silencing of *nt*5 gene led to the destruction of this nucleic acid pathway (p<0.01) (Fig. 2C). However, the growth rate of mutant *S. aureus* Δ*nt*5 was similar with the wild-type *S. aureus* (*S. aureus* WT) (Fig. 2D), which indicated *nt*5 gene had little direct influence on the growth rate of *S. aureus* ATCC 25923.

**Fig. 2.**
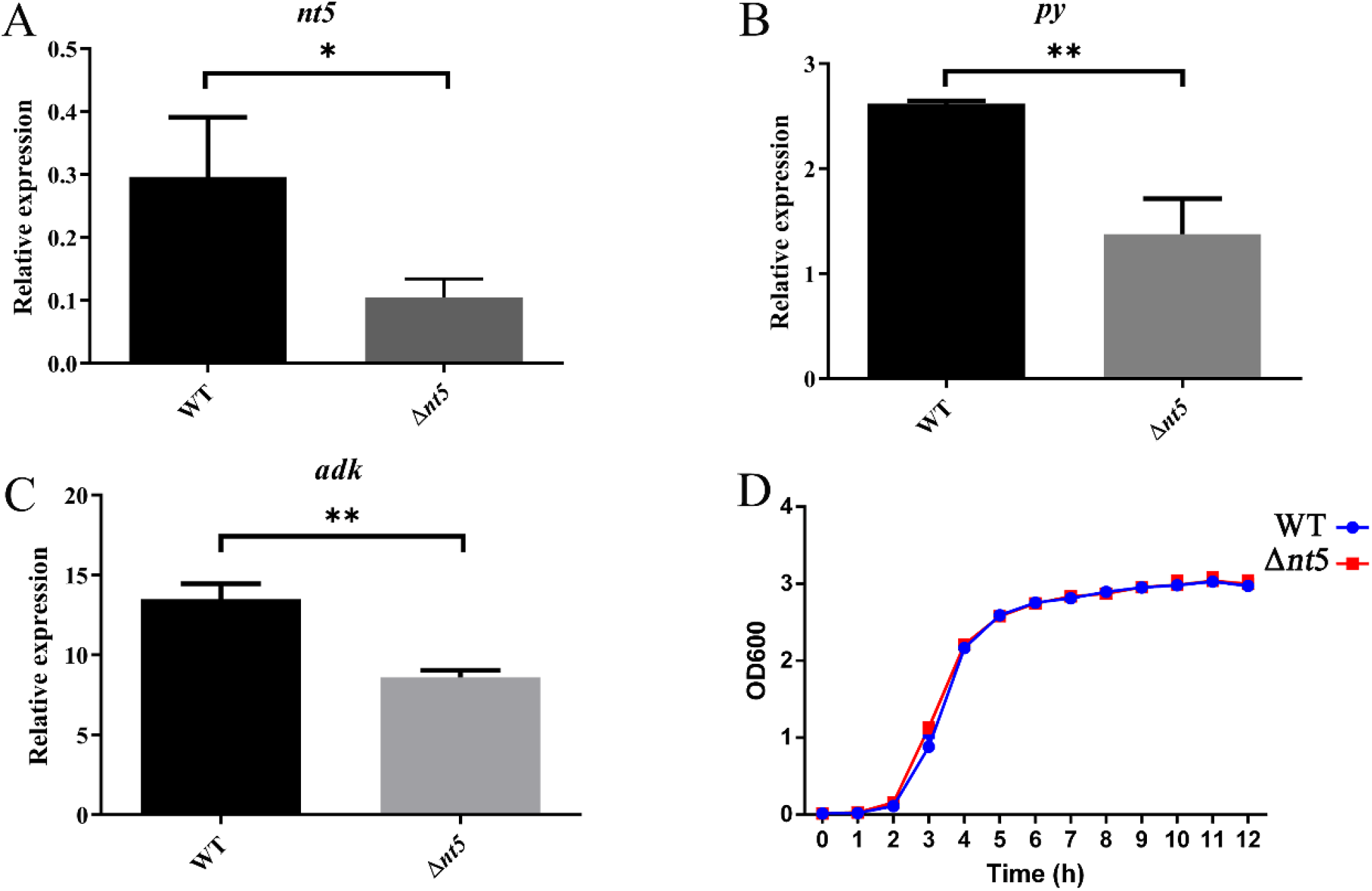
The mRNA expression level of target gene in gene silence mutant Δ*nt5*. (A) The *nt5* gene expression in Δ*nt5* mutant was deceased in comparison with WT. (B) The mRNA level of *py* gene was downregulated in Δ*nt5* compared with WT. (C) Compared with WT, lower *adk* mRNA was detectable in Δ*nt5*. (D) The mutant strain Δ*nt5* exhibited a similar growth rate to the WT. WT: wild type, *py*: pyruvate kinase, *adk*: adenylate kinase, *P<0.5, **p<0.01.

### *nt*5^C166T^ mutant phenotype and relative gene expression changes in response to DAP exposure

We further studied the phenotype changes of the *nt*5^C166T^ mutant Δ*nt*5 upon DAP treatment. The MIC values of DAP on Δ*nt*5 mutant was proximately 0.5 mg/L, which had little significant differences from the WT upon DAP treatment. Upon 0.5 MIC (0.25mg/L) DAP treatment, the mRNA level of *nt*5 was downregulated by 2.90 times in *S. aureus* WT, while there was no significant change of *nt*5^C166T^ in *S. aureus* Δ*nt*5 when compared to without DAP-exposed bacteria (p<0.05) (Fig. 3A). The results indicate *S. aureus nt*5 gene is key and sensitive to antibiotics DAP exposure, which has been confirmed in our previous studies [6]. The mRNA level of *py* was 1.96-times upregulation in WT upon 0.25mg/L DAP exposure, but *py* had no significant change in Δ*nt*5 with DAP treatment (p<0.05) (Fig. 3B). In addition, the mRNA level of *adk* gene was upregulated respectively by 2.06 and 2.35 times in WT and Δ*nt*5 upon DAP exposure (p<0.001, p<0.0001) (Fig. 3C).

**Fig. 3.**
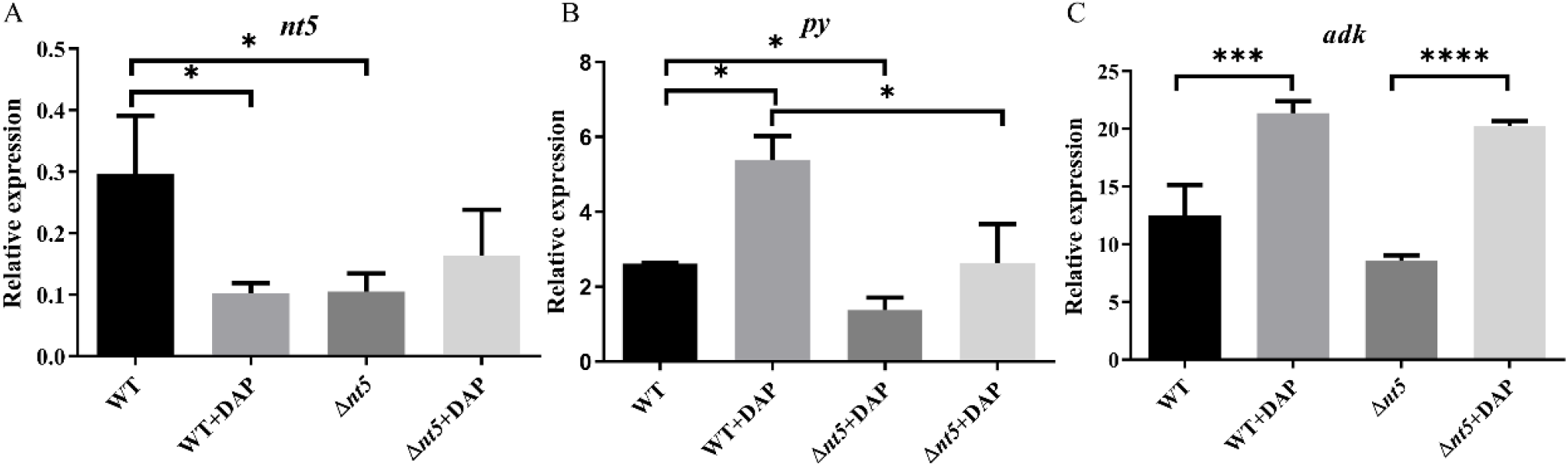
The *nt5*-mediated *py* and *adk* levels in response to DAP exposure in *S. aureus* Δ*nt5*. (A) The *nt5* was decreased in WT upon 0.25 mg/L DAP treatment. (B) The mRNA level of *py* was upregulated by 0.25 mg/L DAP treatment in *S. aureus* WT and Δ*nt5*. (C) The mRNA level of *adk* was upregulated in WT and Δ*nt5* upon 0.25 mg/L DAP treatment. DAP: daptomycin, *P<0.5, ***p<0.001, ****P<0.0001.

Above all, it showed that mutation of the key pathogenic gene *nt*5 would induce gene transcription changes of *nt*5-related genes *py* and *adk* upon DAP exposure. Although that the silence of *nt*5 gene leads to the destruction of nucleic acid pathway, but the silence of *nt*5 gene has no direct influence on growth of bacteria. So far, we speculate that the silence of *nt*5 gene may affect the virulence of bacteria, and we further investigate *nt*5 roles for *S. aureus* pathogenesis, specially focusing on abscess formation.

### *nt*5 deficiency impedes *S. aureus*-infected abscess formation

10^6^ colony-forming units (CFU) of *S. aureus* WT, Δ*nt*5 was respectively inoculated into BALB/c mice by intravenous injection to investigate the role of gene *nt*5 for abscess formation. Ten mice were included for each group of *S. aureus* inoculation, but two mice died of *S. aureus* WT infection before infection experiment ending. Mice were killed after bacterial infection for 5d, and right kidneys were collected for observation of abscess profiles by H&E staining and colony counting respectively.

It was noticed that WT formed more abscesses in mouse kidneys compared with the Δ*nt*5 (Fig. 4A). The abscess lesions were further observed in kidney tissues by H&E staining analysis. There were several obvious lesions surrounded by infiltrating neutrophils in the kidney of mice inoculated with *S. aureus* WT.

**Fig. 4.**
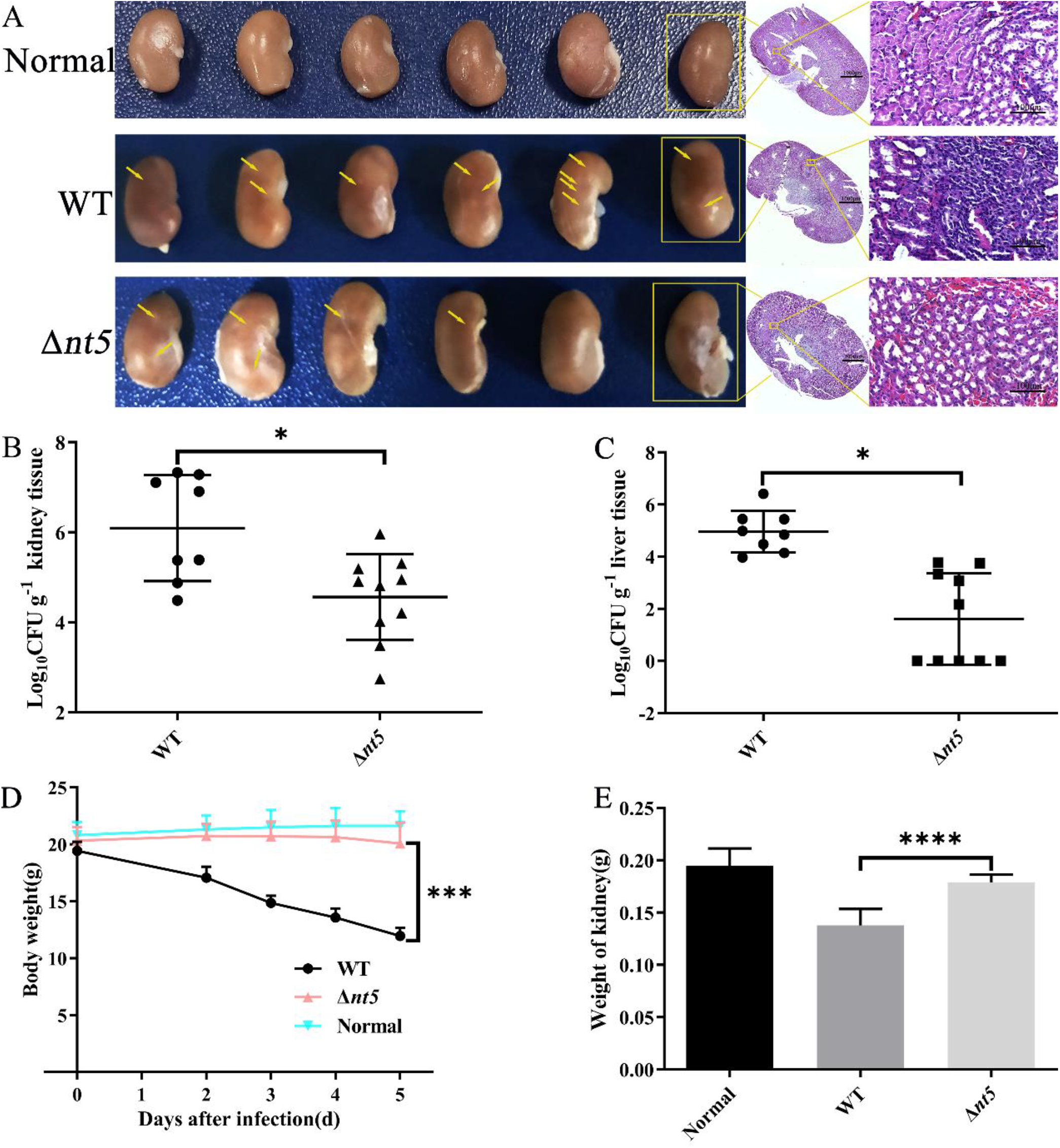
Numbers of kidney abscess and liver bacterial loading were less in *S. aureus* Δ*nt5*-infected mouse than WT infection. (A) *S. aureus* Δ*nt5*-infected kidney abscess. Yellow arrows indicated abscess lesions. (B-C) Staphylococcal loading in mouse kidneys (B) and livers (C) after intravenous injection of 10^6^ CFU of *S. aureus* WT and Δ*nt5* for 5 d. Ten mice were included in each group, but only 8 mice were finally enumerated in the *S. aureus* WT-infected group (n = 8) due to 2 mice died before experiment ending. (D) Body weight of the mice inoculated with Δ*nt5* was increased compared with WT. Normal group: normal BALB/c mice injected with PBS. (E) Mass weight of mouse kidneys infected with *S. aureus* Δ*nt5* was nearly recovered to that of normal mouse. *P<0.5, ***p<0.01,****P<0.0001.

Moreover, 6×10^6^ (CFU/g tissue) *S. aureus* WT bacteria were detectable in mouse apostematic kidney. Under same conditions, but there was 1.5×10^5^ (CFU/g tissue) *S. aureus* Δ*nt*5 surviving in mouse apostematic kidney tissue (Fig. 4B). So, the living *S. aureus* Δ*nt*5 in mouse kidney was about a 40-times amount reduction compared with that of *S. aureus* WT infection (P<0.05), which indicates *nt*5 gene silence greatly impedes infection ability of *S. aureus*.

Similarly, the colony formation quantity of *S. aureus* Δ*nt*5 surviving in mouse liver showed a 300-times reduction as compared with WT (P<0.05) (Fig. 4C), which demonstrated infection ability of Δ*nt*5 to mouse liver was also obviously attenuated.

### *nt*5 silence recovers *S. aureus*-infected mouse body weight

The *S. aureus* Δ*nt*5 and WT strains were assessed to have influence on the body weight of the infected mice. The mouse weight with Δ*nt*5 injection was about 20g, which showed no significant change in body weight compared with the normal BALB/c mice injected with PBS. But the weight of mouse infected with *S. aureus* WT was greatly reduced to a mean value of 13g, which indicated *S. aureus* WT-induced infection led to an obvious decrease of mouse body weight. Compared with *S. aureus* WT, the mouse weight with Δ*nt*5 injection was recovered and increased up to the normal mouse weight (p<0.001) (Fig. 4D). The *nt*5 gene expression attenuates *S. aureus*-infected mouse body weight.

In addition, the weight of each kidney from *S. aureus* Δ*nt*5-infected mouse was about 0.18g, while a kidney *ex vivo* was reduced to 0.14g in *S. aureus* WT-infected mouse (P<0.0001) (Fig. 4E). It was consistent with changes of *S. aureus*–infected mouse body weight. In general, *nt*5 gene contributes to the infection capacity of *S. aureus*, and *nt*5 silence recovers *S. aureus*–infected mouse body weight as similar to the normal mouse without infection.

### *nt5* gene silence promotes phagocytosis to *S. aureus* by macrophages *in vitro*

In order to investigate the mechanism of *nt*5^**C166T**^-weakened abscess formation, we respectively co-cultured *S. aureus* WT or *S. aureus*Δ*nt5* with macrophage cell line RAW264.7 and THP-1 *in vitro* to study *nt5* role in immune response (20). As a result, *S. aureus* Δ*nt5* was more easily phagocytosed by RAW 264.7 and THP-1 cells when co-culture from 15 to 60 min by counting bacterial colonies on TSA at timed intervals (Fig. 5A and 5B). After co-culture for 75-90 min, *S. aureus* WT and Δ*nt5* were almost clear away.

**Fig. 5.**
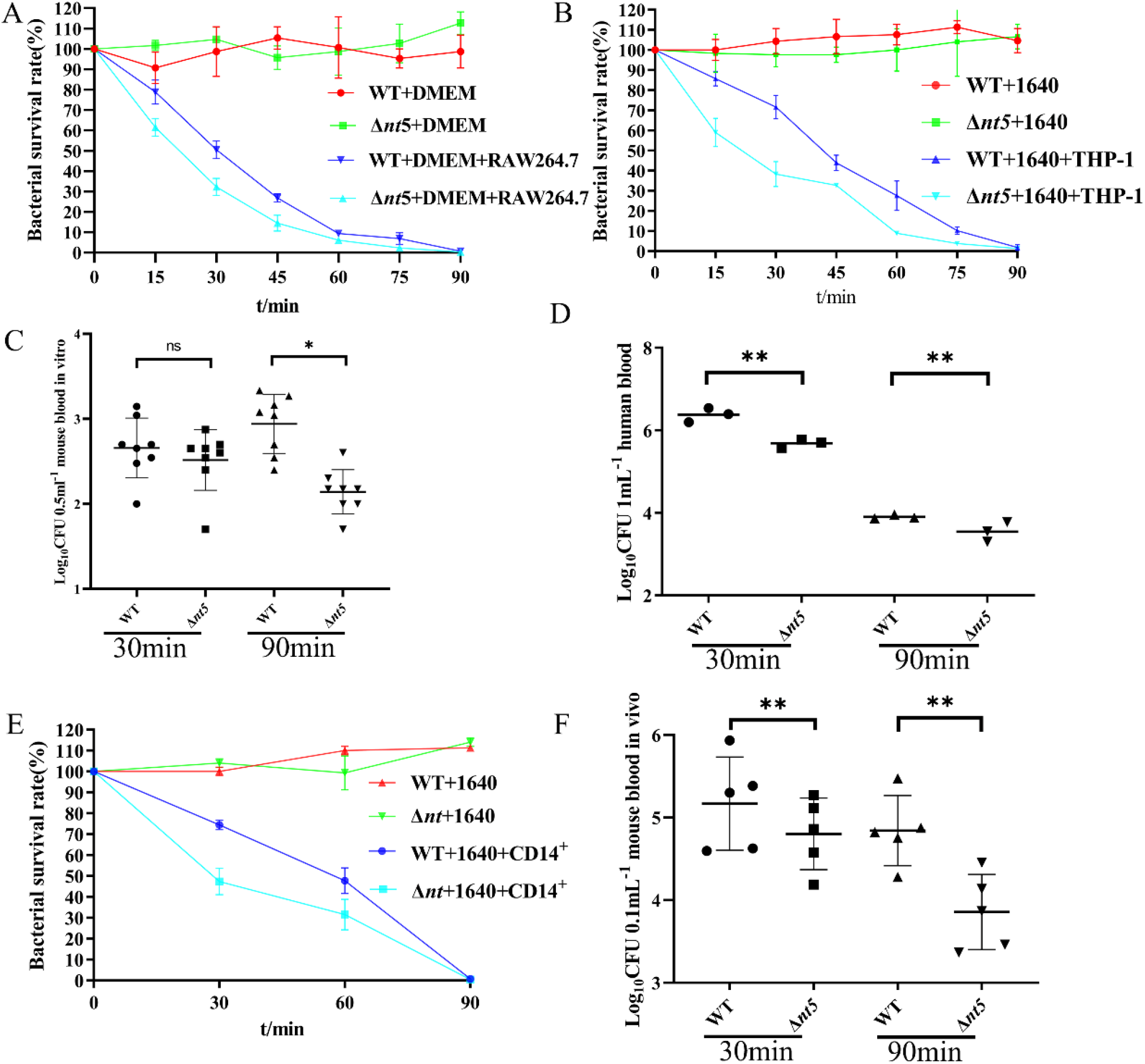
Phagocytosis of *S. aureus* by host immune cells. (A-B) Phagocytosis of *S. aureus* by rat macrophage cell line RAW264.7 (A) and human monocyte cell line THP-1 (B). The survival rate of bacteria was calculated by recording the surviving CFU of *S. aureus* on TSA. (C-D) The surviving ability of *S. aureus* Δ*nt5* in peripheral blood *in vitro. S. aureus* WT, Δ*nt5* was co-cultured *in vitro* with blood sample collected from 8 BALB/c mice (C) and 3 human volunteers (D) by recording bacterial load at timed intervals. (E-F) Phagocytosis of *S. aureus* Δ*nt5* by human (E) and mouse (F) CD14^+^ immune cells *in vitro*. The survival rate of bacteria was calculated by recording the surviving CFU of *S. aureus* on TSA.

In addition, we incubated bacteria with host peripheral blood *in vitro* respectively collected from 8 mice and 3 human volunteers at 37°C for 30, 90 min to examine the survival ability of *S. aureus* WT and Δ*nt5*. There was little difference between WT and Δ*nt5* on bacterial load at incubation with mouse blood for 30 min (Fig. 5C). We speculated the anticoagulant EDTA on the blood anticoagulation tube probably influenced biological activity of immune cells, and EDTA affected release of complement C5a, thus affecting the phagocytosis effect of macrophages (21,22). Under same conditions at incubation with each donor’s blood for 30 min, the remaining bacterial load of Δ*nt5* was decreased than the WT (p<0.01) (Fig. 5D), which indicated *nt5* gene silence improves phagocytosis of macrophages to *S. aureus*. But at incubation for 90 min, the surviving bacterial quantity of the mutant Δ*nt*5 was less than the *S. aureus* WT in both of two groups (p<0.05, p<0.01) (Fig. 5C and 5D), which indicated *S. aureus* Δ*nt*5 displayed a defect in staphylococcal escape from phagocytic killing compared with the *S. aureus* WT.

Human innate immune system firstly responds to *S. aureus* infection, so we further isolated CD14^+^ monocytes from human peripheral blood mononuclear cells (PBMC) to detect the immune response to *S. aureus* Δ*nt*5 exposure. We sorted out CD14^+^ cells from 3 volunteers’ blood samples and co-cultured CD14^+^ cells with *S. aureus* WT and Δ*nt5* respectively. The survival of bacteria was detected by recording bacterial load at timed intervals via the formation of colonies on TSA (Fig. 5E). *S. aureus* Δ*nt5* was more easily cleared out by human CD14^+^ cells, which indicates that *nt5* gene plays an important role in immune escape for *S. aureus* infection. Because CD14^+^ monocyte cells bind to cell wall peptidoglycan of *S. aureus* and then activate phagocytosis to protect human body suffering from *S. aureus* infection (23).

### *nt5* gene is required for bacterial survival in mouse blood

In order to further verify the role of *nt5* gene in host immune response, we directly injected 10^6^ CFU*S. aureus* WT and Δ*nt5* into BALB/c mice to detect the killing effect of host immune system on bacteria. In short time of bacterial infection for 30 and 90 min *in vivo*, the quantity of surviving *S. aureus* WT was higher than the mutant Δ*nt5*, which shows the mutant Δ*nt5* is more easily cleared by mouse immune system (p<0.01) (Fig. 5F).

In conclusion, we have revealed that *nt5* is required for staphylococcal survival in host as an extracellular pathogen, and *S. aureus nt5* gene plays a critical role for staphylococcal escape from the phagocytic killing of host immune defense system to promote their survival in host blood and tissues.

### *nt5* silence has no influence on hemolysis

We detected whether the silence of *nt*5 gene affected the hemolytic activity of bacteria. We inoculated 10^5^ CFU bacteria on blood agar plate and cultured at 37°C for 16 h. The silence of *nt*5 gene had no influence on the hemolytic activity of bacteria (Fig. 6A and 6B). Therefore, the attenuation of bacterial virulence caused by the silence of *nt*5 gene is mainly reflected in the formation of abscess.

**Fig. 6.**
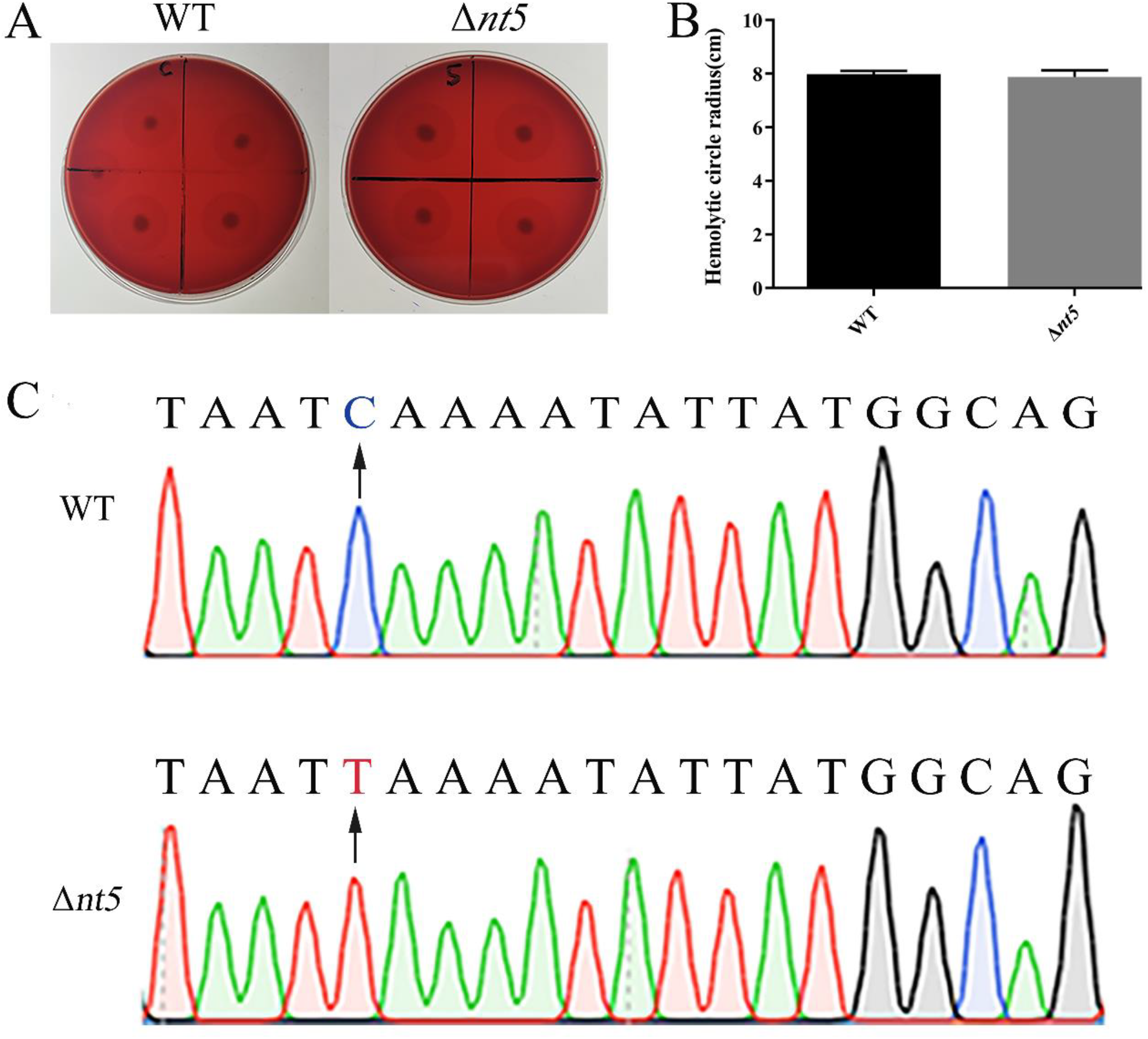
Hemolytic activity of *S. aureus* Δ*nt5*. (A-B) Size of hemolytic ring formed by WT and Δ*nt5* on blood agar plate. (C) The *nt*5 gene sequencing of *S. aureus* Δ*nt5* colony from mouse liver homogenate and kidney homogenate.

In addition, we verified gene stability of *nt*5 silence mutant grown on the TSA plate that was derived from the liver and kidney tissue homogenate extracted from *S. aureus*Δ*nt5*-infected mouse. The DNA sequencing showed the growing bacteria were consistent with *S. aureus* Δ*nt5* (Fig 6C), which indicates the *nt*5^**C166T**^ mutation via pnCasSA–BEC single base editing system is genetically stable.

## Discussion

Almost all archaea and many bacteria achieve adaptive immunity through a diverse set of CRISPR-Cas systems, which has proven to be extraordinarily simple and efficient in DNA and RNA editing, nucleic acid detection, and disease diagnosis (24,25). The CRISPR/Cas9 system has been engineered for gene editing in various organisms which is originally confirmed as a bacterial immune syste (18). The pCasSA system is one kind of specific CRISPR/Cas9 designed for *S. aureus* gene knockdown (18). But most of *S. aureus*, especially clinical isolates, cannot survive due to the double-stranded DNA breaks induced by the pCasSA system and the lack of the homologous recombination repair process. A novel base editing system based on CRISPR RNA-guided cytidine deaminase (17) is available to construct *S. aureus* Δ*nt*5 since this genome editing approach does not require formation of double-stranded DNA breaks or provision of a donor DNA template, which allows for rapid and efficient genetic manipulation and drug-target exploration in *S. aureus* (24).

Base editing method is an efficient gene silence strategy, which induces single-nucleotide changes in the DNA of living cells. There are two kinds of base editors including the cytidine editor and the adenosine editor. Efficient editing by the C/T conversion “base editor” requires the presence of a PAM motif with NGG nucleotide sequence, which has an editable window from positions 4 to 8 in the spacer sequence (18). This PAM requirement limits the number of sites in the bacterial genome. An editable site of the pnCasSA–BEC system is essential for highly efficient base editing. In addition, it has a disadvantage that mutation site could not be verified by mRNA levels compared with CRISPR/Cas9 system. For example, the mRNA expression level of *nt*5 gene in our study is still observed in Δ*nt*5, so DNA sequencing is required to confirm gene mutations into terminate codons (26).

Herein, we focus on physiological changes in *S. aureus* with *nt*5 gene silence. The silence of *nt5* gene influences the expression of *py* and *adk* genes, but it does not affect the growth of bacteria, which indicates metabolic compensation of nucleic acid pathway in bacteria (27). And we have revealed that *nt*5-mediated nucleic acid pathway in *S. aureus* is obviously disturbed due to increase of its pathogenic genes *py* /*adk* and *nt*5 decrease in response to antibiotics treatment. These observations are in line with the bacterial proteome profiling of *S. aureus* ATCC 25923 upon DAP treatment (7). Meanwhile, it also shows that DAP not only disrupts bacterial cell membrane, but also affects the *nt*5-mediated nucleic acid pathway in *S. aureus*, which contributes to the effective antibacterial activity of DAP (28). Therefore, disruption of *nt*5-mediated signaling paves possible therapeutic opportunities for treatment of infections and provides a direction for finding new target of DAP.

*S. aureus* usually causes serious infections, with an annual incidence of 20 to 50 cases per 100,000 people. Approximately 10% to 30% of these patients die from *S. aureus* infection, which accounts for a greater number of deaths than for AIDS, tuberculosis, and viral hepatitis combine (29). Although antibiotics, such as vancomycin and DAP, have powerful antibacterial activity on *S. aureus*, the increase of bacterial drug resistance leads to the gradual failure of antibiotics. It is necessary to explore the pathogenic mechanism of *S. aureus*.

By now, this is the first study to reveal role of *nt*5 gene in abscess formation and immune evasion. The *in vivo* analysis in mouse sepsis model indicates that *S. aureus nt*5 gene is required for kidney abscess formation and is helpful for bacterial evasion from the host immune response during infection. Immune evasion is an important process in which microorganisms directly modulate host-immune system in response to pathogens (30). Of note, in some bacteria, NT5 contributes to immune evasion by dephosphorylating adenosine mono-, di-, or tri-phosphates, thereby increasing the concentration of adenosine.

In *S. aureus*, NT5 activity is the mediator of the virulence attributes of *adsA* (16). And NT5 has classic lipoprotein signal sequence which belongs to the haloacid dehalogenase superfamily and is similar to haemophilus influenzae lipoprotein E. A critical role of NT5 in haemophilus influenzae is the degradation of external riboside (30). *S. aureus* adsA is an adenosine synthase to own NT5 activity. AdsA converts neutrophil extracellular traps to dAdo during infection, and dAdo production represents a general immune evasion mechanism for microbial pathogens (31). Pathogenic bacteria use synthesized dAdo to evade host defenses (14). The production of dAdo by *S. aureus* adsA induces caspase-3-mediated death of human macrophages (14). *Streptococcus suis* produces dAdo by NT5 causing monocytopenia in mouse blood *in vivo*, which suggests the immunosuppressive activity of Ado on immune responses *in vivo* (8). Perturbation of adenosine levels is likely to affect host immune responses during infection (31,32).

Moreover, the global proteome and phosphor-proteome of sepsis kidney injury also reveal that caspase-1 is significantly up-regulated in the cecal ligation and puncture-induced sepsis, which mediates gasdermin D cleavage and activation (33). Gasdermin D dependent pyroptosis cell death has been revealed for the first time in nonseptic acute kidney injury (34). In the kidney abscess caused by *S. aureus*, it’s valuable to further investigate whether does exist a similar mechanism with cecal ligation and puncture-induced sepsis.

*Ex vivo* infection of whole blood is a valuable tool to study *S. aureus* virulence factors and the host innate immune responses. In order to consider effects of cellular mediators, the coagulation cascade must be inhibited to avoid clotting. However, EDTA, a strong chelator of Ca^2+^ as well as Mg^2+^, blocks coagulation as well as complement entirely (21). Blocking of C5a or C5aR1 attenuates phagocytosis and increases the extracellular growth of *S. aureus* in blood (22). Besides the host response, the anticoagulants might directly affect the bacteria growth. EDTA as a chelator of Ca^2+^ is present on the blood tube, which has a strong inhibitory effect on *S. aureus* growth when *S. aureus* is phagocytosed by macrophages *in vitro*. Hence, the citrate or heparin as anticoagulant should be selected to collect host blood sample, which has little effect on host immune response and bacterial growth (21).

*S. aureus* secretes a number of host-injurious toxins, among the most prominent of which is the small β-barrel pore-forming toxin α-hemolysin. Initially named based on its properties as a red blood cell lytic toxin, early studies suggested a far greater complexity of α-hemolysin action as nucleated cells also exhibited distinct responses to intoxication (35). However, our experimental results show that gene *nt5* does not affect the hemolytic activity of *S. aureus*. We speculate that the virulence of gene *nt5* is mainly manifested in sepsis and immune escape.

In conclusion, we have provided direct evidences of the role of *nt*5 gene toward staphylococcal infection ability. We have revealed *nt*5 gene of *S. aureus* is required for host infection and abscess formation. These results are helpful for the development of NT5 inhibitors as a new strategy for rendering *S. aureus* susceptible to human host defence.

## Acknowledgments

This work was supported by the Projects of International Cooperation and Exchanges from Sichuan Province (2020YFH0094), Chengdu Science and Technology Bureau (2020-GH02-00056-HZ).

---

**S1 Table.**
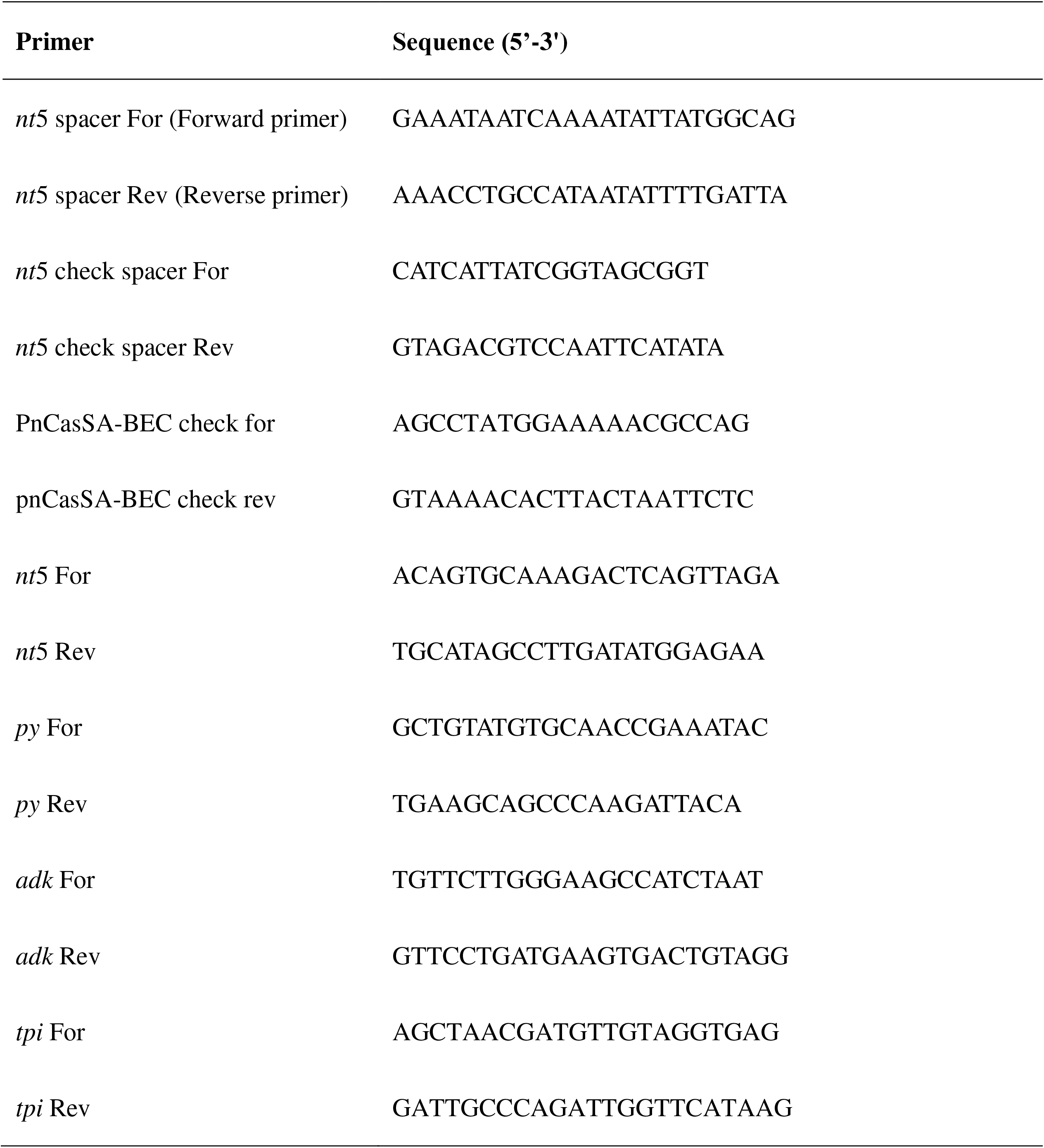
Primers used in this study.

